# Improvement of the memory function of a mutual repression network in a stochastic environment by negative autoregulation

**DOI:** 10.1101/279109

**Authors:** A B M Shamim Ul Hasan, Hiroyuki Kurata, Sebastian Pechmann

## Abstract

**Background:** Cellular memory is a ubiquitous function of biological systems. By generating a sustained response to a transient inductive stimulus, often due to bistability, memory is central to the robust control of many important biological processes. However, our understanding of the origins of cellular memory remains incomplete. Stochastic fluctuations that are inherent to most biological systems have been shown to hamper memory function. Yet, how stochasticity changes the behavior of genetic circuits is generally not clear from a deterministic analysis of the network alone. Here, we apply deterministic rate equations, stochastic simulations, and theoretical analyses of Fokker-Planck equations to investigate how intrinsic noise affects the memory function in a mutual repression network.

**Results:** We find that the addition of negative autoregulation improves the persistence of memory in a small gene regulatory network by reducing stochastic fluctuations. Our theoretical analyses reveal that this improved memory function stems from an increased stability of the steady states of the system. Moreover, we show how the tuning of critical network parameters can further enhance memory.

**Conclusions:** Our work illuminates the power of stochastic and theoretical approaches to understanding biological circuits, and the importance of considering stochasticity to designing synthetic circuits with memory function.

## Background

Memory is ubiquitous in biological systems [1–4]. Characterized by a continued response to a transient stimulus [5], cellular memory has been found to aid in the robust control of diverse biological functions such as synaptic plasticity [6], differentiatio n [7, 8], cell cycle transition [9], or gene regulation [10]. Memory in cellular circuitry is tightly linked to bistability, i.e. the presence of two stable steady states [11, 12]. To this end, memory is achieved if an input signal evokes a switch to an alternative steady state where the system remains over time even after the input signal has disappeared [11, 12].

Our general understanding of design principles of biological networks has improved dramatically during recent years [13]. Feedback loops as elementary components provide common control mechanisms of cellular networks [14]. For instance, negative feedback can facilitate adaptation and oscillation [15], while positive feedback loops, which play pivotal roles in cellular signaling [16], often promote signal amplification, bistable switches [17] and memory [11]. Indeed, a common circuit architecture that is known to give rise to memory is based on interlinked positive feedback network loops [18, 19]. Several experimental and theoretical analyses of such network architectures that can give rise to memory have been reported [7, 12, 20, 21, 22–25]. These studies have revealed many general properties underlying successful memory, for instance, that ultrasensitivity may be sufficient to generate two stable states [26], and that the transition periods between the bistable states compose an important characteristic of a cellular memory module [19]. In all cases, bistability and memory can use a transient input signal to yield a robust cellular system [27, 28]. This property is directly visible from the characteristic hysteresis behavior observed for bistability and memory systems. Experimental examples confirm these findings, such as the presence of hysteresis behavior and bistability in the budding yeast galactose-signaling networks [18]. Based on these general principles that link network architecture to systems behavior, the blossoming field of synthetic biology has contributed many useful designs of genetic circuits based on a quantitative understanding of biological networks [29–31] with myriad implications for biotechnology, biocomputing and gene therapy [13, 26, 32]. However, the rational design of a robust memory function can be implemented via many alternative mechanisms that remain incompletely understood [21, 29].

Importantly, the vast majority of genes in a cell is normally only expressed at very low levels, thus giving rise to substantial stochastic fluctuations [33–39]. As a result, the analysis and design of biological circuits cannot be based solely on deterministic properties of the network topology alone. In response, much progress has been made in better understanding the role of stochasticity to network function [40–44]. For instance, careful work has delineated the sources of intrinsic noise in small transcriptional networks [45] and how noise may propagate through network architectures [38, 40], as well as how feedback loops regulate bistability [41, 46] and intrinsic noise [47]. Similarly, extensive work has established general principles underlying the design of genetic circuits that exhibit bistability and memory [11, 41, 42, 43, 44].

Recent examples of opposing behavior further highlight the importance of considering the contribution of stochasticity to cellular circuitry. On the one hand, it has been shown that noise can induce multimodality and even stochastic memory in a system that, according to a deterministic description, lacks bistability [48, 49]. On the other hand, stochastic fluctuations in gene expression levels often reduce or disrupt the memory function of biological networks [21, 50, 51]. Equally importantly, seminal technical and conceptual advances have made strong progress in the efficient and accurate approximation of multivariate nonlinear stochastic systems [52, 53].

Here, we investigate how a negative autoregulation network architecture can improve the sustained memory function of a mutual repression network in a stochastic environment. Our work extends the previously published regulated mutual repression network (MRN) [12] to the regulated mutual repression network with negative autoregulation (MRN-NA) to investigate the effect of a negative autoregulation loop on memory function. The network architecture is characterized by a mutual repression cycle that can give rise to bistability and memory, adopted from two well-characterized mutual repression networks, the system consisting of LacI and TetR in *E. coli* that displays a bistable gene expression memory module [24], and the mutual repression of the two repressors cI and Cro that yield a bistable memory module in the bacteriophage *λ* [21].

To explore and compare the memory behavior of MRN-NA network model, as well as how it compares to the MRN model, deterministic rate equations, stochastic simulations and theoretical analyses of Fokker-Planck equations were employed to identify principles of the robustness of the memory function. We demonstrate how negative autoregulation can reduce intrinsic noise and thus improve memory function by increasing the stability of the steady states. Negative autoregulation can for instance be achieved by the ability of proteins to bind and sequester their own mRNA. Systematic analyses of successful memory as a function of central model parameters that describe the mutual repression cycle highlight principles and limits of memory function in these mutual repression networks. Through systematically comparison of these model systems under stochastic fluctuations our results contribute important insights into the functioning of biological circuits in the presence of noise. Moreover, our work highlights the importance of considering stochasticity when designing synthetic circuits with memory function.

## Results

To investigate the effect of negative autoregulation and stochasticity on network memory function, we constructed a model network following a previously established graphical notation [54, 55] (Fig. 1). Specifically, we modelled a mutual repression network with negative autoregulation (MRN-NA) that extends our previously published mutual repression network (MRN) [12]. The network consists of proteins *y*(1), *y*(2) and *y*(3). An input signal *S* induces the synthesis of *y*(1), which activates and represses the syntheses of *y*(2) and *y*(3), respectively. The syntheses of *y*(2) and *y*(3) are mutually repressed with cooperativity. Moreover, negative autoregulation controls the synthesis reactions of *y*(2) and *y*(3) (Fig. 1). The addition of negative autoregulation permits to investigate how negative feedback loops may affect memory. While modulation of several parameter values should tune the presence of memory, the present analysis focuses on a direct back-to-back comparison of two network architectures that only differ in the addition of negative autoregulation circuits. To compare the memory regions of MRN-NA and MRN networks across our deterministic, stochastic, and theoretical analyses, the values of the corresponding kinetic parameters as well as the steady state levels of *y*(2) and *y*(3) were conserved as much as possible in discrete parameter space ([12], Table 1, Texts S1 and S2) (see Methods).

**Table 1.** List of kinetic parameters used in the two gene regulatory networks

| **Kinetic parameters** | **Definition** |
| --- | --- |
| $k(1), k(3), k(4), k(6), k(7)$ | protein synthesis rate constants |
| $k(2), k(5), k(8)$ | degradation rate constants |
| $K(1), K(2), K(3), K(4)$ | dissociation constants of activators / repressor |
| $k(9), k(10)$ | protein synthesis rate constants of negative autoregulation |
| $K(5), K(6)$ | dissociation constants of negative autoregulation |
| $n$ | Hill coefficient |

**Fig 1.**
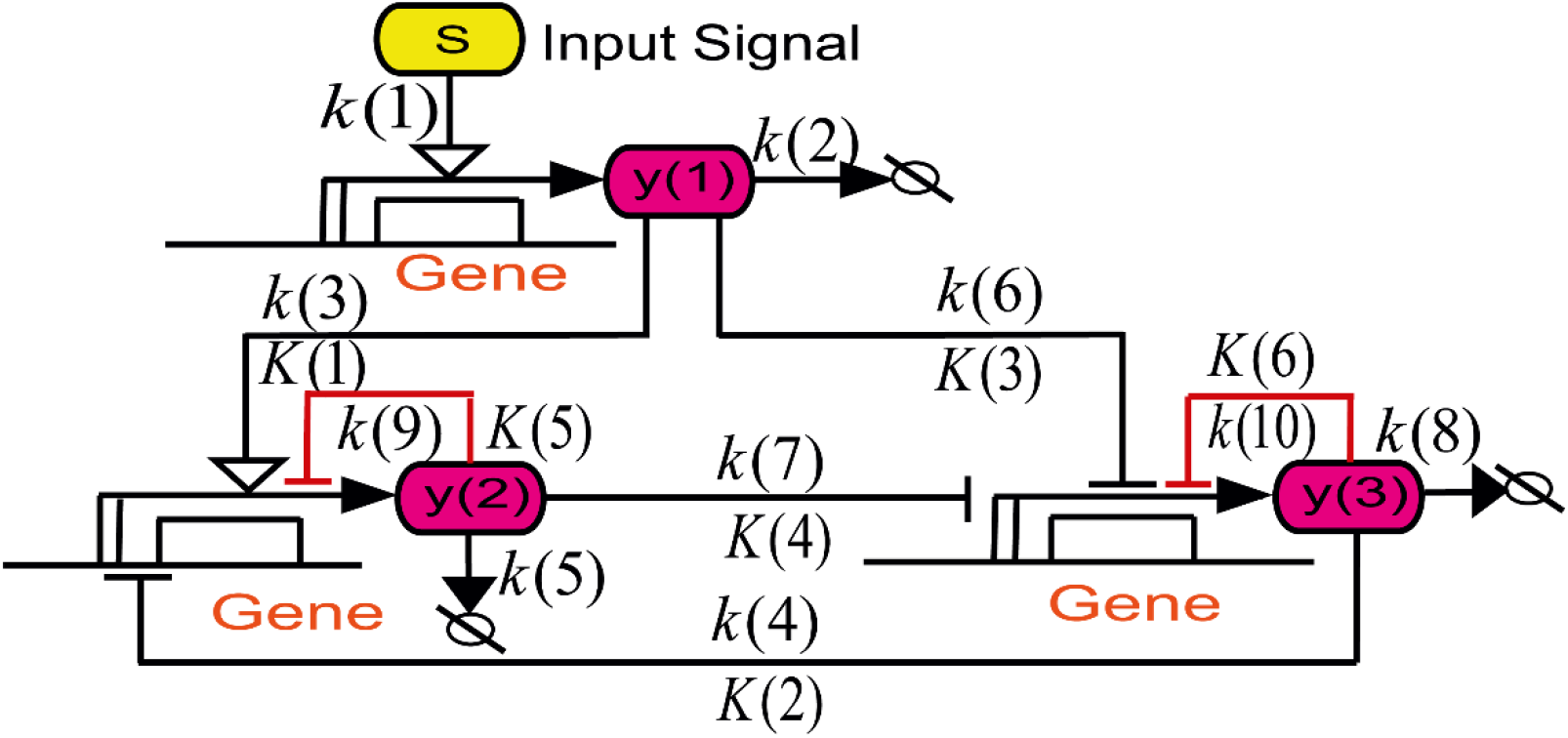
Schematic of the mutual repression network models with negative autoregulation. Wiring diagram of the mutual repression network with negative autoregulation (MRN - NA) model. The added negative autoregulation reactions that regulate the protein synthesis of *y*(2) and *y*(3) are highlighted in red. All reaction rate constants *k* and dissociation constants *K* are listed in Tables 1 and 2. In this work, the MRN-NA is compared to the previously published MRN [12] that is identical with the exception of the negative autoregulation loops.

### MRN-NA model exhibits robust of memory

Similar to our previously reported results on the memory function of the MRN [12], the MRN-NA network exhibits robust memory functionality. The deterministic solution to the time evolution of the MRN-NA model showed persistent memory after the end of the period during which the signal *S* was applied for both Hill coefficients of *n* = 7 and *n* = 8 (Fig. 2A). Strong hysteresis behavior was observed for both *y*(2) and *y*(3) levels as a function of increasing and decreasing *S* (Fig. 2BC, Fig. S1). Stochastic simulations of the MRN-NA model as described by the birth and death processes of Eqs. (1–3) revealed strong and persistent memory both with Hill coefficients *n* = 7 (Fig. 2D) and *n* = 8 (Fig. 2E). Importantly, the MRN alone could not sustain memory for either *y*(2) or *y*(3) at *n* = 7 in a stochastic context [12], but the MRN-NA yielded strong and sustained memory due to the addition of negative feedback loops (Fig. 2D and Fig. S3, S5).

**Table 2.** Settings of the kinetic parameters for the two models

|  | MRN | MRN-NA |
| --- | --- | --- |
| Common parameters<br>between the MRN and<br>MRN-NA | $k(1) = 100$<br>$k(2) = 1$<br>$k(3) = k(6) = 18.1$<br>$k(4) = 61.23 > k(7),$<br>$k(7) = 43.1$<br>$k(5) = k(8) = 0.8$<br>$K(1) = K(3) = 9$<br>$K(2) = K(4) = 43$<br>$n = 8$ | $k(1) = 100$<br>$k(2) = 1$<br>$k(3) = k(6) = 18.1$<br>$k(4) = 61.23 > k(7),$<br>$k(7) = 43.1$<br>$k(5) = k(8) = 0.8$<br>$K(1) = K(3) = 9$<br>$K(2) = K(4) = 43$<br>$n = 8$ |
| Specific parameters to<br>the MRN-NA | | $k(9) = k(10) = 4.1$<br>$K(5) = K(6) = 9$ |

**Fig 2.**
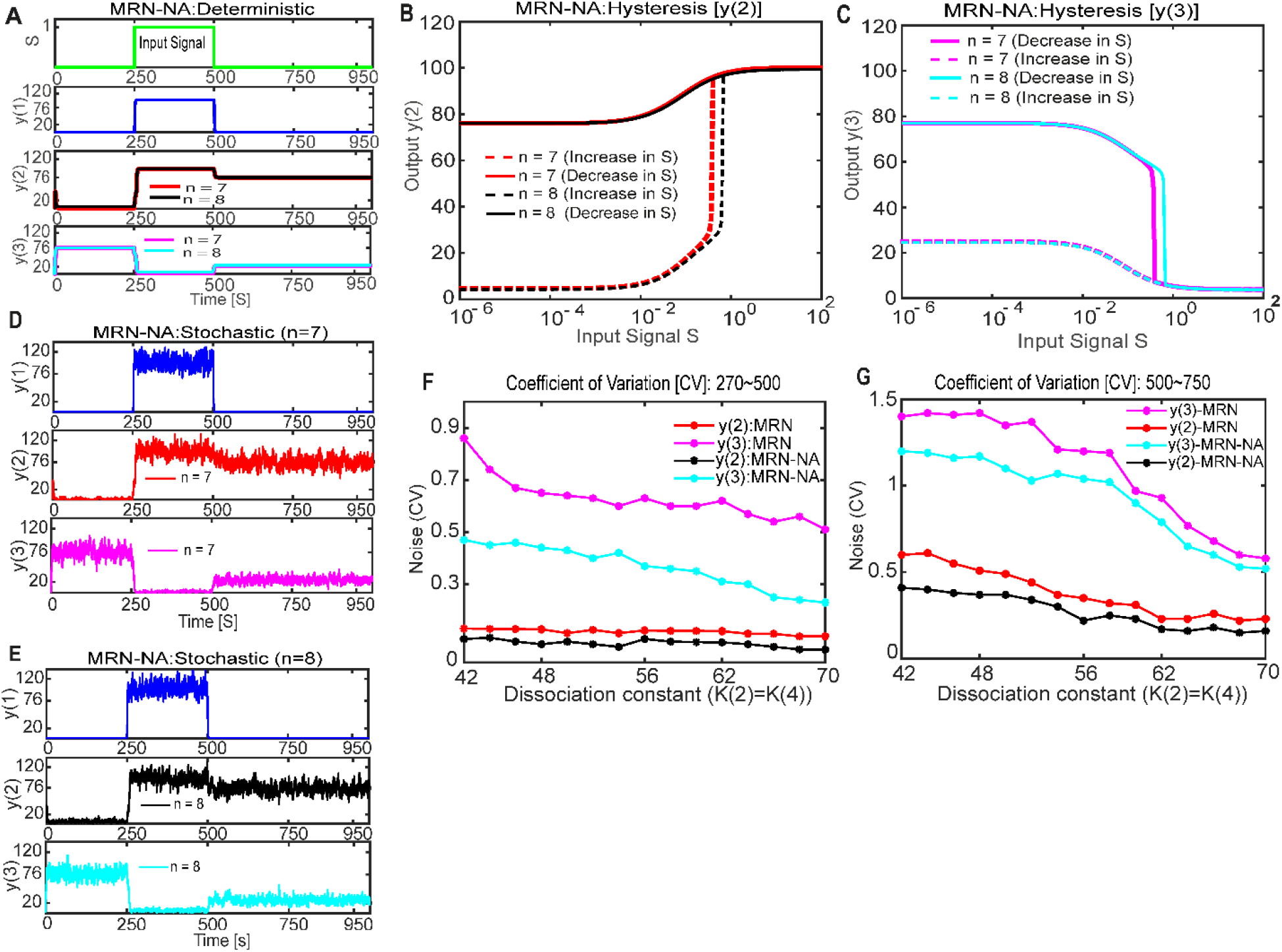
Deterministic and stochastic simulations of the MRN-NA model. (A) Deterministic simulation of proteins *y*(1), *y*(2) and *y*(3) at different Hill coefficients. The input signal *S* is applied from simulation steps 250 to 500. The dissociation constants are set to *K*(2) = *K*(4) = 43; all other corresponding parameter values are set as the same as the previously published MRN [12] (Table 2). The simulated time evolution is shown for *y*(2) and *y*(3) at *n* = 7 (red and magenta lines), and at *n* = 8 (black and cyan lines). (B) The hysteresis curves of *y*(2) at different Hill coefficients *n* = 7 and *n* = 8 are consistent with the numerical integration of the rate equations (C) The hysteresis curves of *y*(3) at different Hill coefficients *n* = 7 and *n* = 8 are consistent with the numerical integration of the rate equations. (D) Trajectories of the stochastic simulation of *y*(1), *y*(2) and *y*(3) at Hill coefficient *n* = 7. (E) Trajectories of the stochastic simulation of *y*(1), *y*(2) and *y*(3) at Hill coefficient *n* = 8. (F,G) Stochastic fluctuations in the MRN and MRN-NA models during the interval from simulation step 270 to 500 (F) and during the period from simulation steps 500 to 750 (G). The coefficients of variation (CVs) are computed from the simulated stochastic trajectories during the signal period as a function of changing dissociation constants *K*(2) = *K*(4) at Hill coefficient *n* = 8. For more optimal comparability between the two models, the parameters associated to the negative autoregulation reactions were tuned to conserve the high steady state levels between both models. Shown are the CVs of *y*(2) (MRN: red line; MRN-NA: black line), and *y*(3) (MRN: magenta line; MRN-NA: cyan line).

To further understand how the addition of negative autoregulation increases the robustness of the memory function, we sought to quantify and compare the intrinsic noise in the protein levels of *y*(2) and *y*(3) in the MRN and MRN-NA models. It is well established that noise or a stochastic perturbation can flip gene expression levels from one to the other steady state [39, 45]. To obtain a reliable and comparable estimate, we computed the coefficients of variation (CVs) of *y*(2) and *y*(3) during the active signal *S* period; the interval from simulation step 270 to 500 was chosen to allow the system to respond to the stimulus for 20 simulation steps. Our analysis of noise in protein levels *y*(2) and *y*(3) as a function of the dissociation constants *K*(2) = *K*(4) confirmed a systematic reduction of stochasticity in *y*(2) and *y*(3) levels upon the addition of negative autoregulation (Fig. 2F). While maintaining the same steady state levels, noise was dramatically reduced for *y*(3) (Fig. 2F). Of note, the lower steady state *y*(3) is more susceptible to noise. Similarly, intrinsic fluctuations in *y*(2) during the signal period were reduced in the MRN-NA compared to the MRN model in our stochastic simulations irrespective of the dissociation constants tested (Fig. 2F). Even more important to the stability of the steady states are the intrinsic fluctuations after the end of the signal period. The levels of intrinsic noise are overall slightly higher, likely because the signal does not stabilize gene expression levels any longer. However, our corresponding analysis reveals that the negative autoregulation reduces, as expected, variability in the system also after the stop of the signal (Fig. 2G and Fig. S4). Taken together, our results suggest that the negative autoregulation played a vital role in increasing the persistence of stochastic memory by reducing intrinsic noise.

### Negative autoregulation enhances the memory region

Our analyses show that the MRN-NA model can exhibit memory, which however critically depends on persistent mutual repression with strong cooperativity. We next set out to systematically map its memory region as a function of Hill coefficient and dissociation constants *K*(2) = *K*(4) in comparison to the previously reported MRN [12] (Fig. 3). Successful memory was defined as sustained levels of *y*(2) and *y*(3) at or near their levels during the applied *S* after the end of the input signal *S* at simulation step 500 until the end of the simulations at simulation step 1000. Moreover, for direct comparison, the kinetic parameter *k*(7) was adjusted to conserve the steady-state levels between the two models (Text S1, [12]). Indeed, the high steady-state levels of both the models were set to the same expression values, while the low steady-state levels were tuned be as similar as possible. Memory was assessed in both the deterministic and stochastic context (see Methods). Accordingly, two memory regions were defined: deterministic memory and stochastic memory (Fig. 3).

**Fig. 3.**
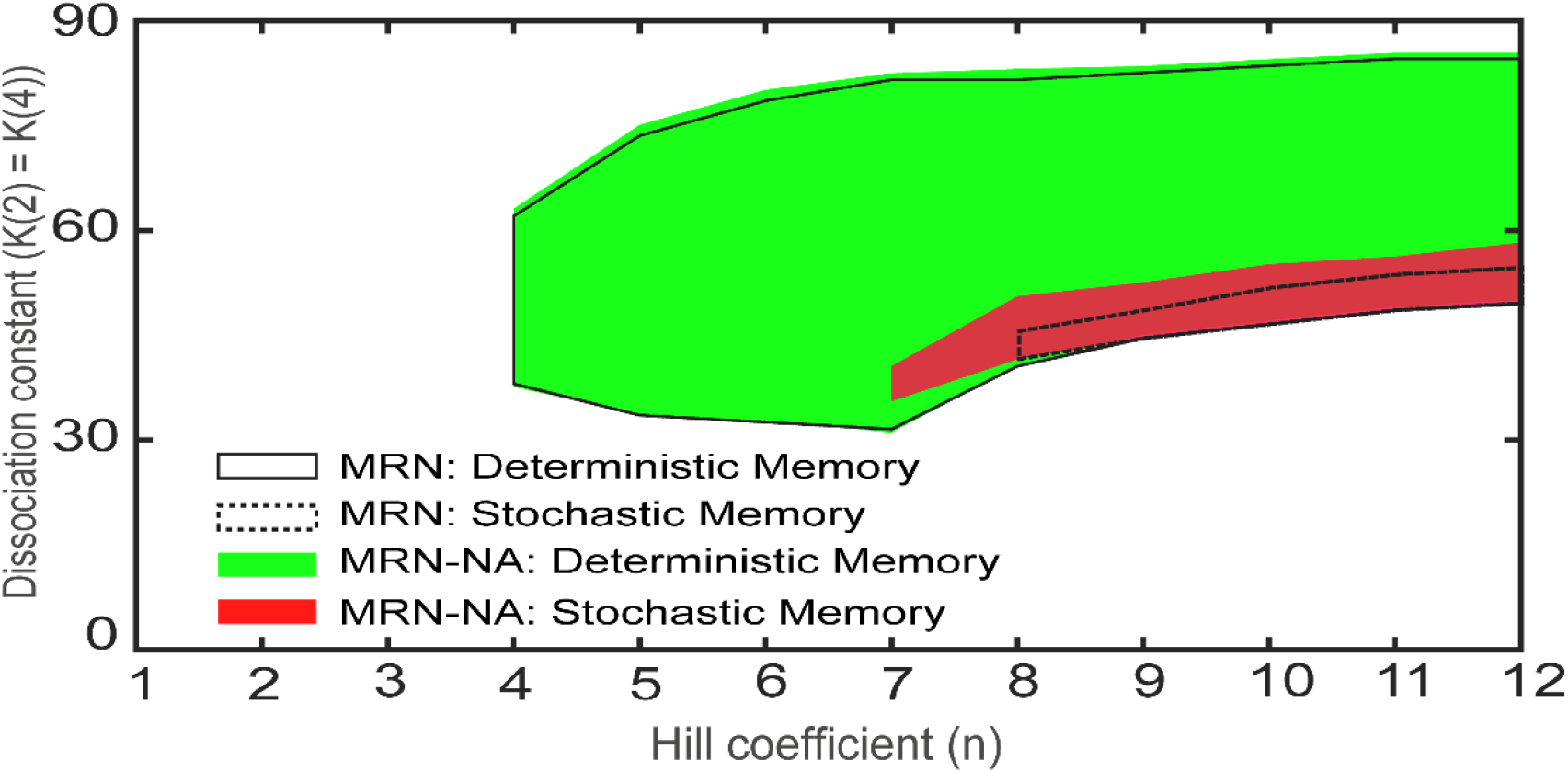
Phase diagram of the memory regions for the MRN and the MRN-NA models. Comparison of the memory regions of the MRN and MRN-NA models. The deterministic (solid line) and stochastic (dotted line) memory regions of the MRN are shown together with the deterministic (green area) and stochastic (red area) memory regions of the MRN-NA as function of the Hill coefficient and dissociation constants.

In both the MRN and MRN-NA models, deterministic memory could be observed for Hill coefficients of *n* = 4 or greater (Fig. 3). In contrast, stochastic memory not only required higher Hill coefficients, but also posed different limits on the MRN and MRN-NA models. Without negative autoregulation that suppresses noise, stochastic memory in the MRN model required a Hill coefficient of *n* = 8 or greater (Fig. 3). In the presence of negative autoregulation that reduces noise in protein levels, stochastic memory in the MRN-NA model could be observed from *n* = 7 (Fig. 3). Thus, the negative autoregulation relaxes the need for strong cooperativity to generate stochastic memory.

The stochastic memory region for the MRN-NA model was found to be larger than for the MRN model (Fig. 3, red area), indicating that negative autoregulation of the MRN-NA enhances memory function by reducing intrinsic noise (Fig. 2F,G). While the deterministic memory regions in both networks are essentially the almost same (Fig. 3, green area), it was clearly more difficult to maintain a memory effect under noise in the MRN than the MRN-NA (Fig. 3; red area). The large discrepancy between the deterministic and stochastic memory areas indicate the general challenge of maintaining memory function in the context of stochasticity.

### Stochastic potential of the MRN-NA

Prerequisite for memory is normally the presence of bistability in the system. Thus, a limited bistable regime could explain why such high levels of cooperativity in the mutual repression between *y*(2) and *y*(3) were required to yield memory. To map out the bistable regimes of the MRN-NA in comparison to the MRN [12], theoretical analyses based on the chemical master equations were employed to support the numerical simulation results.

First, the corresponding Fokker-Planck equations (Eqs. (9–13)) describing the time evolution of the probability densities for *y*(2) and *y*(3) levels were derived from the three rate equations (Eqs. (1–3)) [see Methods]. We made use of the quasi-steady-state assumption to derive a one-variable equation for each system (Text S2, [12]). While there are inherent limitations to it [50], the quasi-steady-state assumption is widely used [46]. Since we focus on the steady-state at *t* →∞, applying the quasi-steady-state assumption to *y*(2) or *y*(3) is mathematically reasonable. By using the Fokker-Planck equations, we next estimated the two-dimensional stochastic bistable regions, characterized through the presence of a double well potential, for the MRN and MRN-NA models as a function of the Hill coefficient and dissociation constants (Fig. 4, Fig. S2; Texts S1, S2, S3 and [12]).

**Fig. 4.**
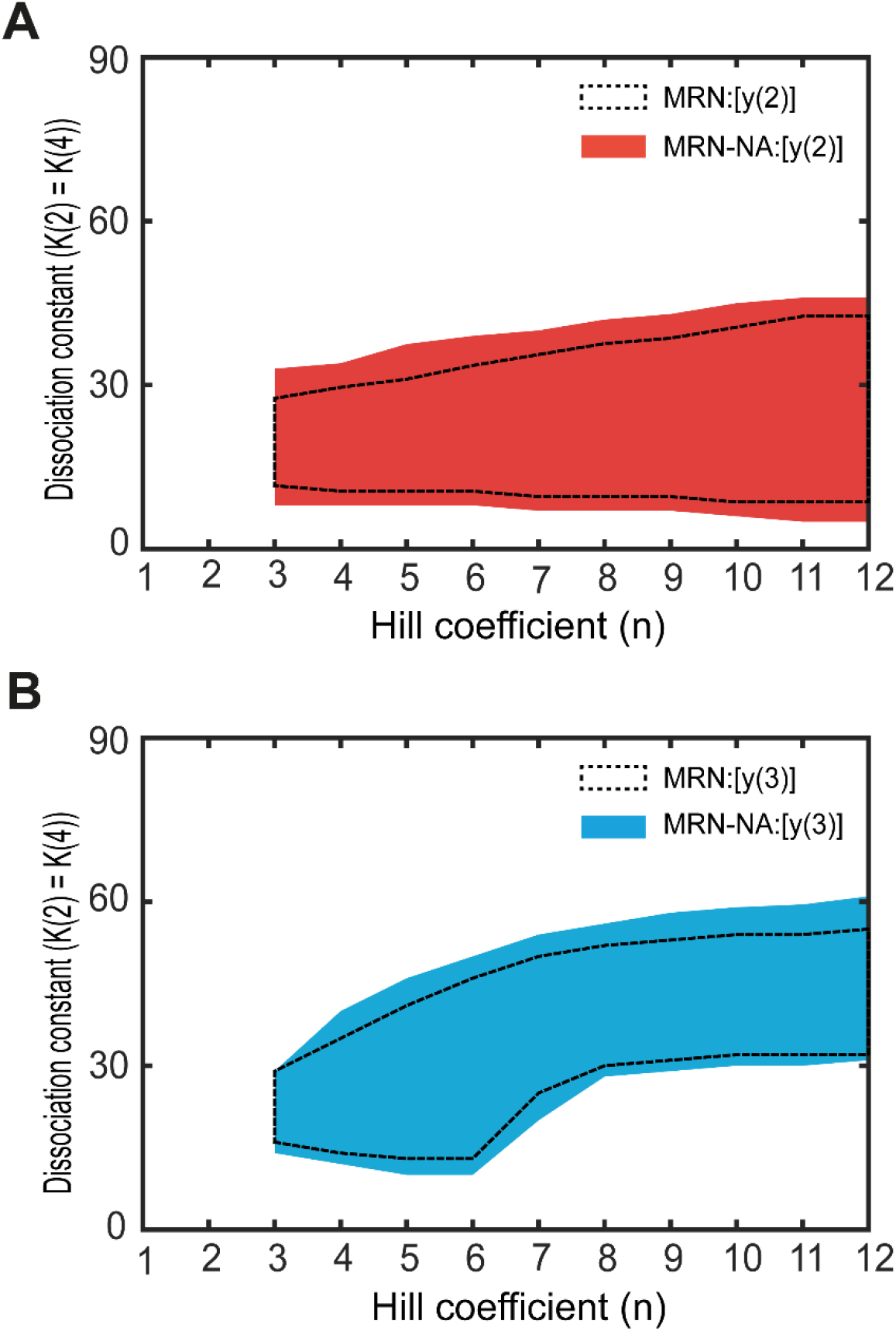
Phase diagram of the stochastic bistable regions. Phase diagrams of the stochastic bistable regions as a function of Hill coefficient and dissociation constant in the mutual repression cycle for *y*(2) in the MRN (dotted line area) and MRN-NA (red area) (A), and for *y*(3) in the MRN (dotted line area) and MRN-NA (cyan area) (B) models.

In these theoretical analyses, both the MRN and the MRN-NA models permitted bistability starting from a Hill coefficient of *n* = 3 for both *y*(2) and *y*(3) (Fig. 4 A, B). However, the bistable regions in the MRN-NA were larger than those in the MRN model (Fig. 4 A, B). To display bistability in a stochastic context, the MRN model required dissociation constants *K*(2) = *K*(4) within a more constrained parameter space (Fig. 4 A, B). These analyses established that the MRN-NA more readily exhibits stochastic bistability than the MRN. Conversely, a limited bistability regime in the MRN in a stochastic context (Fig. 4 A, B) contributes to explaining why this network had a smaller stochastic memory region (Fig. 3).

### Mean first-passage time of the MRN-NA model

Having established that the presence of memory in the MRN-NA model depended on both bistability and robustness of the steady states to stochasticity, we next sought to further explore the origins and limits of memory functionality in context of fluctuating protein levels. To this end, the stability of a steady state of a stochastic system can be estimated by the mean first-passage time (MFPT). The MFPT quantifies the average number of simulation steps it takes for a system to leave a favorable steady state due to random events. As such, the MFPT provides a useful description of the time-scale on which a phase transition is likely to happen [56–58]. Thus, because the presence of a stochastic bistable region only indicates the bimodality of gene expression as prerequisite for memory but not the presence of memory itself, a MFPT analysis can be helpful for identifying the precise conditions under which memory can be attained in a stochastic context.

Here, MFPT analyses were performed between the two stable steady states under the quasi-steady-state assumption (Fig. S2, [12]) (see Methods). Specifically, we calculated the MFPTs of *y*(2) and *y*(3) for the MRN-NA model in comparison to the MRN [12] (Fig. 5); the lower and upper steady-states of *y*(2) and *y*(3) are denoted as 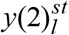, 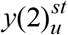, 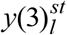 and 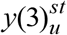, respectively. Similarly, the MFPTs of leaving the lower (*T*_*L*_) and upper (*T*_*U*_) steady states were calculated as a function of the Hill coefficient and dissociation constants in the mutual repression cycle for *y*(2) and *y*(3).

**Fig. 5.**
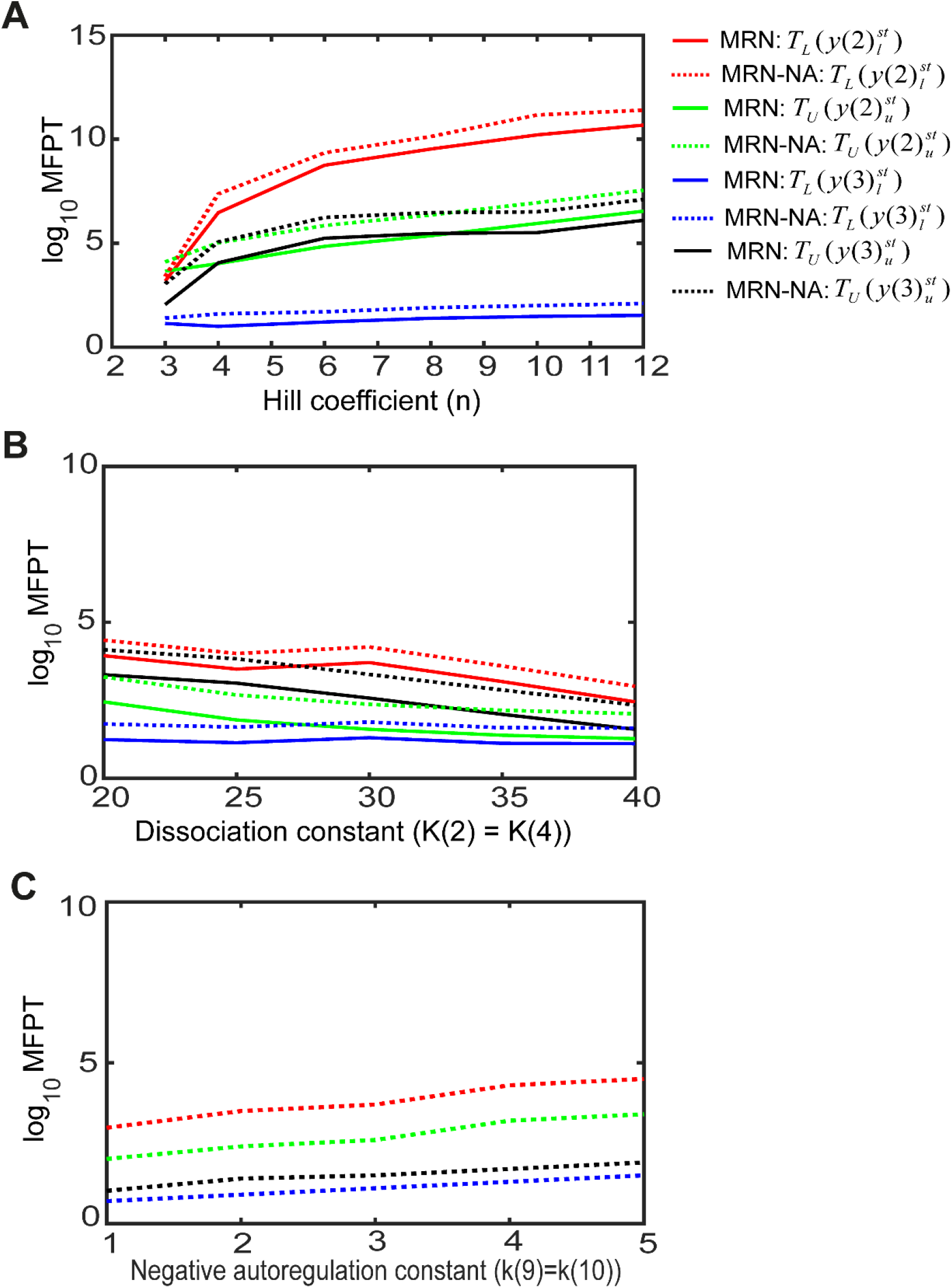
Mean first passage time (MFPT) analysis. (A) The logarithmic MFPTs of the lower and upper steady states of *y*(2) (*T*_*L*_and *T*_*U*_) and *y*(3) (*T*_*L*_ and *T*_*U*_) are shown for the MRN and MRN-NA models as a function of the Hill coefficient *n* at dissociation constants *K*(2) = *K*(4) = 15. (B) The logarithmic MFPTs of the lower and upper steady states of *y*(2) and *y*(3) denoted as above are shown for the MRN and MRN-NA as a function of the dissociation constant*K*(2) = *K*(4) at the Hill coefficient *n* = 3. (C) The logarithm ic MFPTs of the lower and upper steady states of *y*(2) and *y*(3) denoted as above are shown for the MRN-NA as a function of the negative autoregulation constants *k*(9) = *k*(10) at the dissociation constants *K*(2) = *K*(4) = 15 and Hill coefficient *n* = 3.

Due to the asymmetry in the both MRN and MRN-NA models (Table 2), the MFPTs for *y*(2) and *y*(3) showed opposing behavior, as expected (Fig. 5). While the MFPT for *y*(2) to leave the lower steady state, 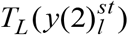, is consistently longer than the corresponding time to leave the upper steady state, 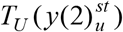, the system is more likely to leave the lower steady state of *y*(3) as indicated by 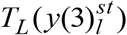 being much smaller than 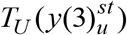(Fig. 5).

An increase in the Hill coefficient as measure of cooperativity exacerbated this trend observed for the MFPTs of *y*(2) and *y*(3) (Fig. 5A). The prolonged MFPTs as a result of increasing Hill coefficients explained why the robustness of sustaining stochastic memory improved with increasing cooperativity. For the parameter space explored, the lower steady state of *y*(2) was more persistent in both models for any Hill coefficients, as evident by 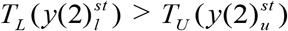(Fig. 5A). Moreover, all MFPTs for the MRN-NA,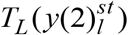,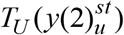,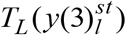, and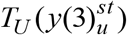, were observed to be longer than their equivalents for the MRN model, for any Hill coefficient (Fig. 5A). These results confirm an increased persistence of the steady states upon introduction of negative autoregulation in MRN-NA compared to MRN, which explains the improved memory function in the MRN-NA system. Specifically, the residence times at the upper and lower steady states of *y*(2) and *y*(3) were extended to achieve the persistent memory after the stop of the signal (Fig. 3D in [12], Fig. 2D).

Next, we investigated the effect of the dissociation constants in the mutual repression cycle on the MFPTs of *y*(2) and *y*(3) (Fig. 5B). Consistently, the MFPTs for all steady states 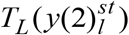, 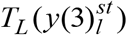, 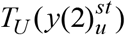 and 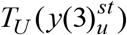,were found to be longer in the MRN-NA than the MRN for any dissociation constants (Fig. 5B). For both models, all MFPTs, 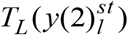, 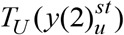, 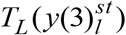 and 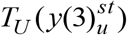 gradually decreased with an increase in the dissociation constant, suggesting that strong binding in the mutual repression cycle is necessary for persistent memory. We also studied the MFPTs for the MRN-NA as a function of the negative autoregulation constants *k*(9) = *k*(10) (Fig. 5C). The MFPTs for the MRN-NA at all steady states, 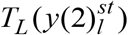, 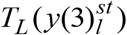, 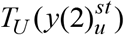 and 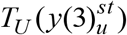, were found to increase with increasing strength of the negative autoregulation. In summary, these MFPT analyses illuminated in detail how memory in the mutual repression networks improved with increasing stability of the steady states as a function of cooperativity or binding strength.

### Probability densities of the steady-state levels in the MRN-NA

The previous analyses revealed a strong dependency of sustained memory on the robustness of the steady-states. To formally establish the probability densities associated with populating the upper and lower steady states of *y*(2) and *y*(3), we calculated the probability densities from the Fokker-Planck Equations (Eq. (15)) as a function of the Hill coefficient and dissociation constants (Fig. 6). The probability density of the upper steady state of *y*(2) increased while that of the corresponding lower steady state decreased with an increase in the Hill coefficient for both the MRN and the MRN-NA models (Fig. 6A). The addition of negative autoregulation in the MRN-NA system visibly made the upper steady state level of *y*(2) more dominant (Fig. 6A).

**Fig. 6.**
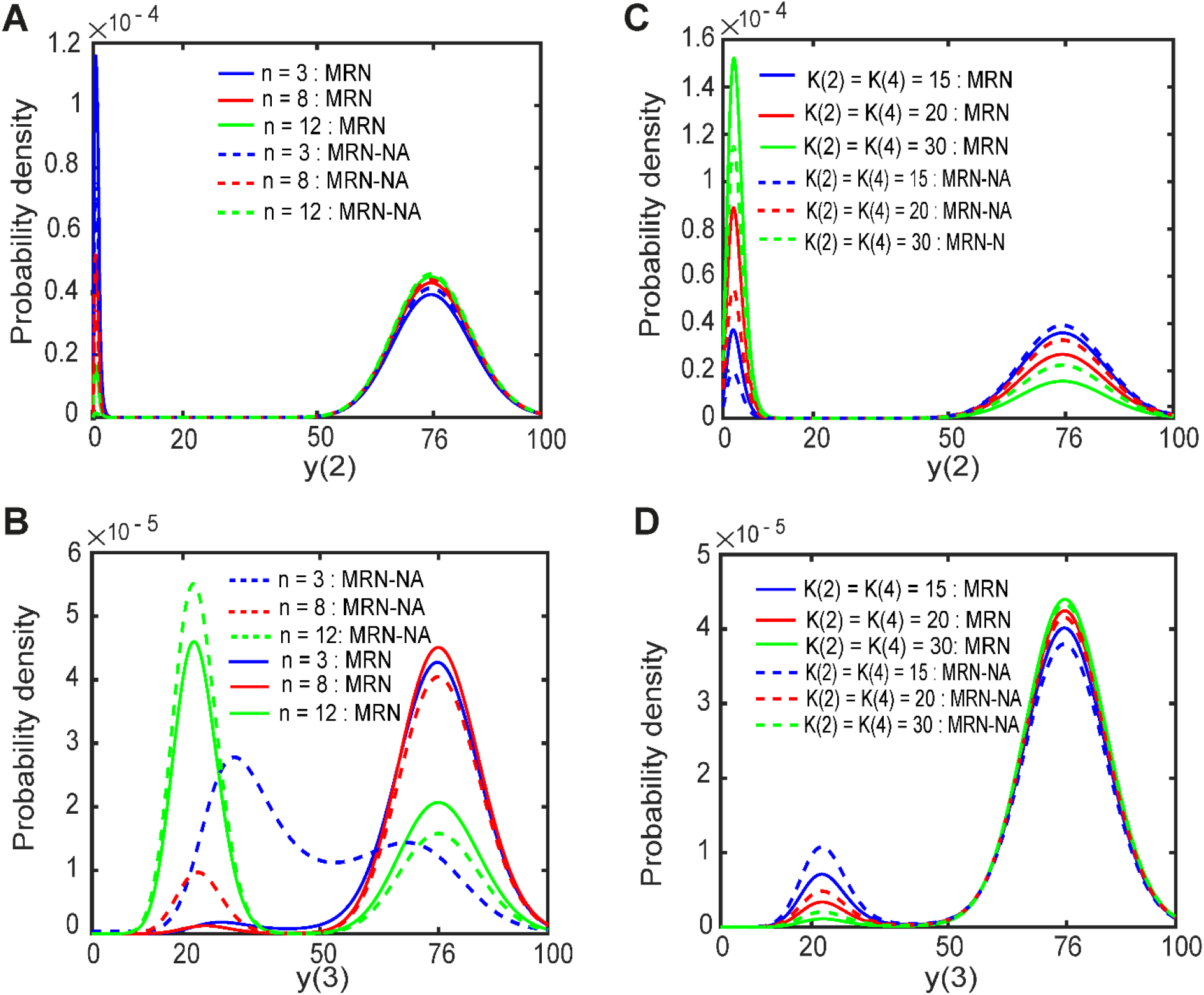
Probability density of the steady-state levels. (A) Probability density of *y*(2) in the MRN and the MRN-NA as a function of the Hill coefficient. (B) Probability density of *y*(3) in the MRN and the MRN-NA as a funct io n of the Hill coefficient at the dissociation constant *K*(2) = *K*(4) = 12 (A, B). (C) Probability density of *y*(2) in the MRN and the MRN-NA as a function of the dissociation constant. (D) Probability density of *y*(3) in the MRN and the MRN-NA as a function of the dissociation constant at the Hill coefficient *n* = 3 (C,D).

The probability density of *y*(3) clearly illustrated the evolution of bistability and subsequent increased probability densities of the lower steady states with increasing Hill coefficient (Fig. 6B). At low Hill coefficient (*n* = 3) the MRN lacked a pronounced probability density of a lower steady state (Fig. 6B, solid blue line) while in the MRN-NA the probability density did not separate well the peaks at lower and upper levels of *y*(3) (Fig. 6B, dashed blue line). This result of weak bistability was consistent with our previous analysis of the bistability regions (Fig. 4). In turn, higher Hill coefficients of *n* = 8 and *n* = 12 promoted first the emergence and then separation of two clearly populated steady states (Fig. 6B). While the upper steady state of *y*(3) was dominant for at *n* = 12 the system switched to a dominant lower steady state for *y*(3) (Fig. 6B). Moreover, the addition of negative autoregulation strongly promoted the population of the lower steady state at all Hill coefficients analyzed (Fig. 6B, dashed lines). By buffering intrinsic noise, this analysis thus confirmed that the negative feedback loops stabilized particularly the low expression states of *y*(3), i.e. those that are most susceptible to fluctuations.

Similarly, the probability densities of *y*(2) and *y*(3) were found to quantitatively and qualitatively also depend on the dissociation constants *K*(2) and *K*(4) in the mutual repression cycle (Fig. 6C,D). At low *K*(2) = *K*(4) = 15, the upper steady state of *y*(2) was dominant in the MRN and MRN-NA models with slightly higher probability density for the MRN-NA model (Fig. 6C). In general, with decreasing values of *K*(2) and *K*(4) the upper steady states became more pronounced in both models, even more so in the MRN-NA (Fig. 6C). In turn, the probability density of the lower steady state of *y*(3) became dominant with decreasing the dissociation constants in both the models (Fig. 6D). Moreover, lower values for *K*(2) and *K*(4) as well as negative autoregulation favored more pronounced bistability (Fig. 6D). The MRN-NA model increased the probability density of the upper steady-state of *y*(2) and the lower steady-state of *y*(3) more than the MRN model. This result can support the persistent memory of the upper steady-state of *y*(2) and lower steady-state of *y*(3) after the stop of the signal (Fig. 6D). Taken together, these results established the theoretical basis for our observation that the MRN-NA model displays an improved memory function in a stochastic context.

### Consistency between the stochastic simulation and the Fokker-Planck equation

To verify the quasi-steady-state assumption for our case, we validated that the solutions to the Fokker-Planck equations were consistent with the probability densities obtained by stochastic simulation of the full system (Figs. 7, 8). The Fokker-Planck equations provided almost the same probability density as the Gillespie stochastic simulation (Figs. 7, 8). We examined the comparison of the numerical and theoretical simulation method in both normal and log scale (Fig. 8). It is shown that lower steady states (first peaks) are quickly transitioning to the upper steady state than second peaks (Fig. 8). This is a sampling issue because initially there are low concentrations of molecules, which can easily move to the upper steady state.

**Fig. 7.**
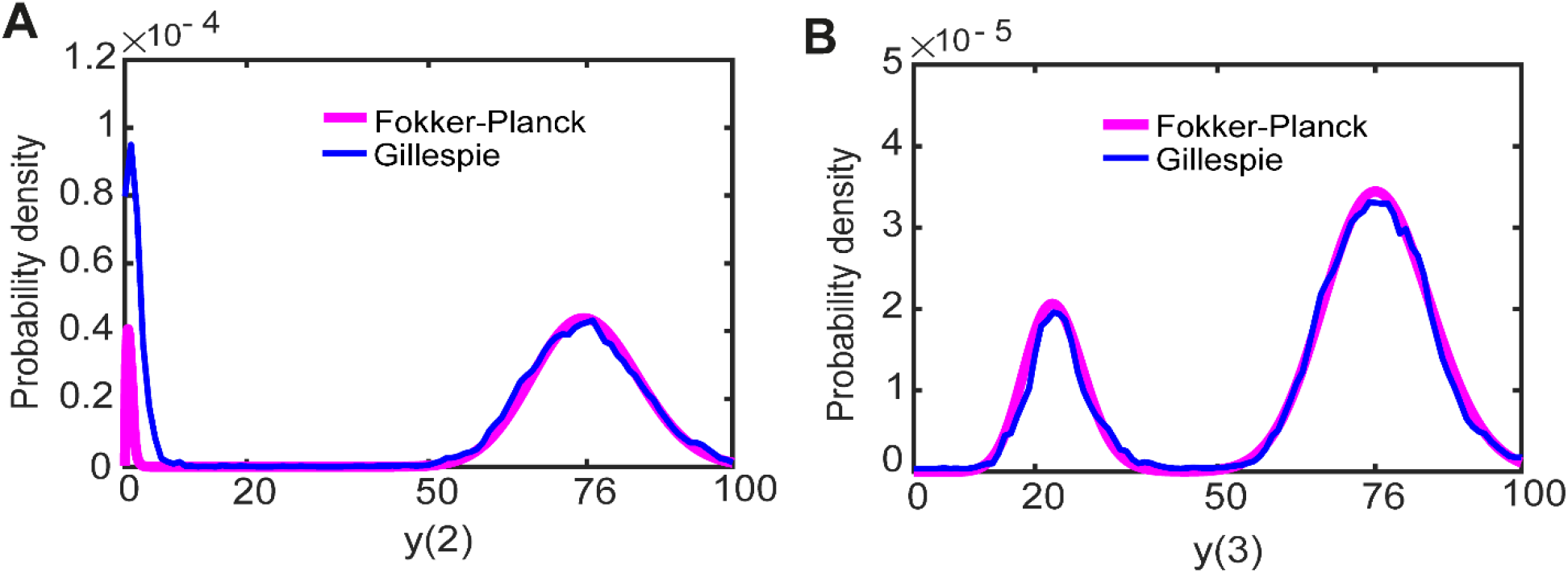
Probability density profile of the MRN-NA model. Probability densities are computed from the Fokker-Planck equations of the MRN-NA model. (A, B) Probability density of *y*(2) (A) and of *y*(3) (B). The parameters are given as *S* = 0, *k*(1) = 100, *k*(2) = 1, *K*(1) = *K*(3) = 9, *K*(2) = *K*(4) = 43, *K*(5) = *K*(6) = 9, *k*(3) = *k*(6) = 18.1, *k*(9) = *k*(10) = 4.1, *k*(4) = 61.04 > *k*(7) = 43.1, *k*(5) = *k*(8) = 0.8, *n* = 8 in the MRN-NA model.

**Fig. 8.**
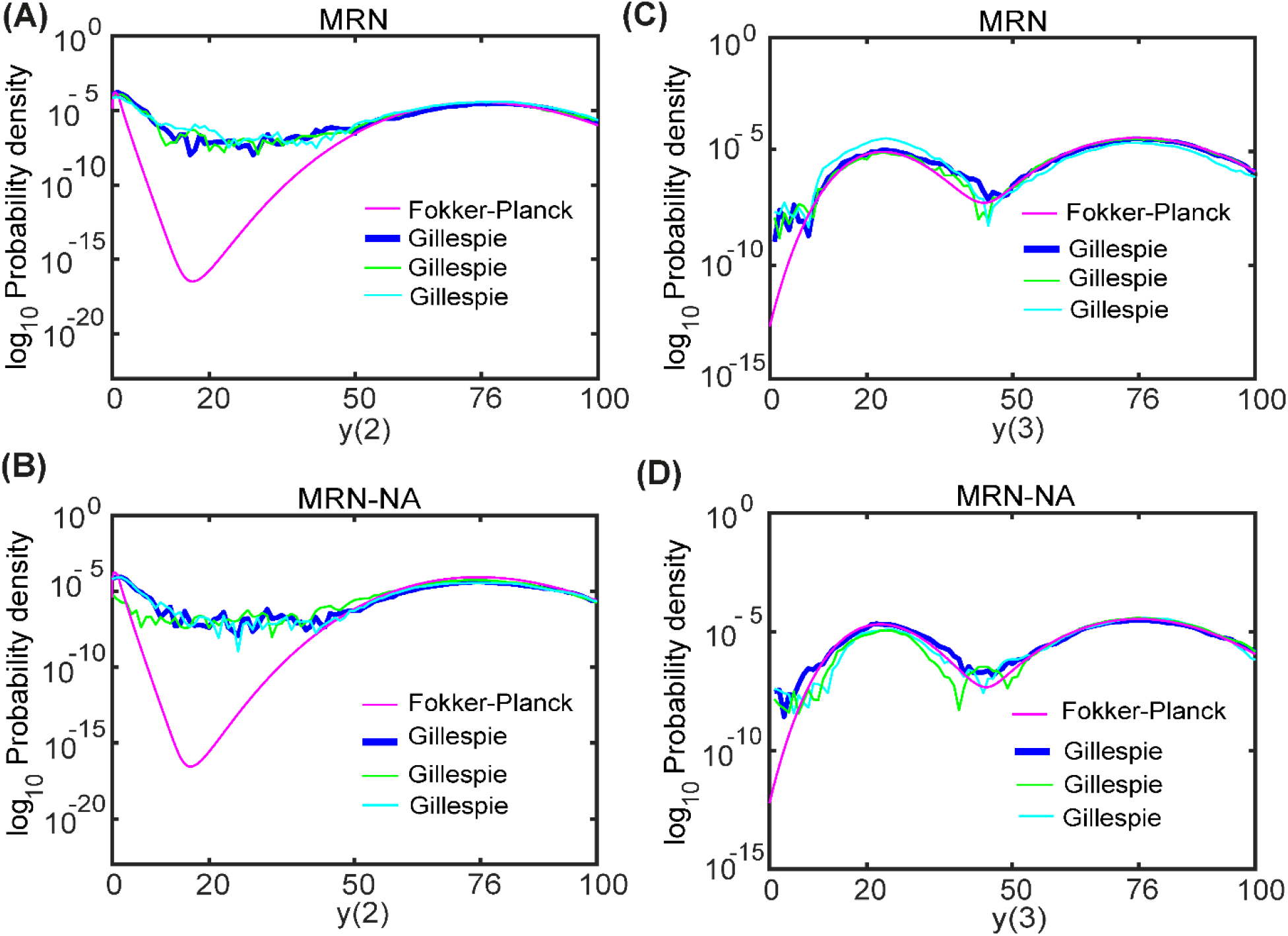
Probability density profile of the MRN and MRN-NA models in log space. Probability densities are computed from the Fokker-Planck equations of the MRN and MRN-NA models. (A, C) Probability density of *y*(2) (A) and of *y*(3) (C) in the MRN model. The parameters are given as *S* = 0, *k*(1) = 100,*k*(2) = 1, *K*(1) = *K*(3) = 9, *K*(2) = *K*(4) = 43, *k*(3) = *k*(6) = 18.1, *k*(4) = 61.23 > *k*(7) = 43.1, *k*(5) = *k*(8) = 0.8, *n* = 8. (B, D) Probability density of *y*(2) (B) and of *y*(3) (D) in the MRN-NA model. The parameters are given as *S* = 0,*k*(1) = 100,*k*(2) = 1,*K*(1) = *K*(3) = 9, *K*(2) = *K*(4) = 43, *K*(5) = *K*(6) = 9, *k*(3) = *k*(6) = 18.1, *k*(9) = *k*(10) = 4.1, *k*(4) = 61.04 > *k*(7) = 43.1, *k*(5) = *k*(8) = 0.8, *n* = 8. For stochastic simulations, three independent trajectories are shown.

## Discussion

By applying numerical integration of the rate equations, stochastic simulations and theoretical analysis of the Fokker-Planck equations, we here investigated the mechanism by which negative autoregulation can improve the memory function in a mutual repression network. Previous work had established the regulated mutual activation network (MAN) and mutual repression network (MRN) as good model systems to study fundamental properties of cellular memory as encoded in genetic circuitry [12]. Here, we systematically decoupled contributions of additional negative autoregulation, as well as strength of cooperativity and dissociation constants in the mutual repression cycle, to the persistence of memory function.

Stochasticity decreased the memory in both models. The MRN-NA however achieved robust, persistent memory in both the deterministic and stochastic approaches at lower cooperativity *n* = 7. Our results suggest that the robustness of the stochastic memory of the MRN-NA could be further improved by increasing the binding strength of the repressor proteins. Moreover, they highlighted the need for finely-tuned parameters combinations to achieve robust memory, as evident from the small stochastic memory regions.

In the present work, defined and validated sets of kinetic parameters were used instead of exhaustive searches for alternative parameter combinations that also give rise to memory. To generalize our findings, phase diagrams of the memory region were derived as a function of the two most critical and previously identified parameters [12]. By doing so, we were able to reveal fundamental insights into the determinants underlying robust stochastic memory. Future work will expand how greater variability in parameter choices, for instance reflecting additional external cellular regulation of constituent genes, may allow to sustain stochastic memory.

Despite accurately simulating the time evolution of protein concentrations in stochastic systems, the Gillespie algorithm is known for its limited power in characterizing sustained memory in more complex networks due to its computational cost. Many time - consuming simulations are required to determine reliably if a model exhibits persistent stochastic memory or not. These shortcomings can be overcome with theoretical approaches. The quasi-steady state assumption of analyzing steady states separately is widely used, performs in general well and is reasonably justified in the presented work. Notably, very recent seminal conceptual and technical advances have provided advanced approximation methods that now start to make even multivariate and nonlinear chemical masters equations and related Fokker-Planck equations amenable to theoretical analyses. For instance the recently introduced linear-mapping approximation converts a nonlinear system to a linear problem via a mean-field approximation [53]. Multivariate systems still become very rapidly too complex for fully analytical solutions. However, these new techniques will vastly improve accuracy and computational efficiency for future analyses of stochasticity in networks.

Equally important for a better understanding of cellular memory is the study of networks of increased complexity that more directly reflect the underlying biology. Known biological examples readily hint at such increased complexity that awaits to be further characterized. For instance, a Notch-Delta mutual repression network serves to communicate between neighboring cells [59] where an increase in Notch activity within a cell decreases that in its neighboring cell. The Notch-Delta mutual repression provides inhomogeneous or opposite protein synthesis in homogeneous cell populations. To capture this phenomenon would require to consider spatial changes in gene expression. Finally, recent papers have outlined many fundamental principles of how to achieve bistability in small networks [60, 61, 62]. The present work here extends these findings by a comparative network analysis that allows to delineate the effect of adding negative autoregulation on cellular memory.

## Conclusions

The addition of negative autoregulation comes at the cost of increased model complexity but clearly improves the robustness of the memory function of a small gene regulatory network in a stochastic context. While the current work has only explored one possible network modification that increases memory, exploration of additional parameter combinations will likely expand our understanding of trade-offs in attaining memory. In summary, we have shown that the addition of negative autoregulation can reduce intrinsic noise and generate persistent stochastic memory. Our mathemat ic a l comparison and theoretical analysis of the two networks contributes to an improved understanding of how genetic circuits can encode biological function and may aid the rational engineering of memory networks for applications in synthetic biology, medicine and biotechnology.

## Methods

### MRN-NA model

We constructed a simple model of a gene regulatory network that consists of two genes encoding a transcription factor that we visualized according to the graphical notation [54, 55] (Fig. 1). Our mutual repression network with negative autoregulation (MRN-NA) consists of proteins *y*(1), *y*(2) and *y*(3). The input signal *S* induces the synthesis of *y*(1), which in turn activates the synthesis of *y*(2) and represses production of *y*(3). Furthermore, the synthesis reactions of both of *y*(2) and *y*(3) are mutually repressed with cooperativity, negative autoregulation governs synthesis reactions of *y*(2) and *y*(3). This system can be bistable and exhibits memory. In formulating our model, we made deliberate use of Michaelis-Menten approximations under the assumption that substrate concentrations are in excess and association equilibria quickly attained. The model is described by the following equations (1-3):

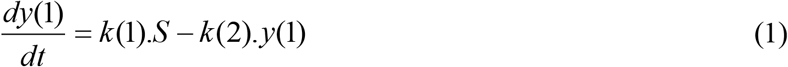

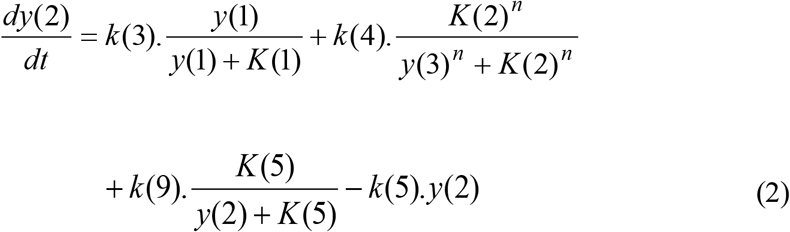

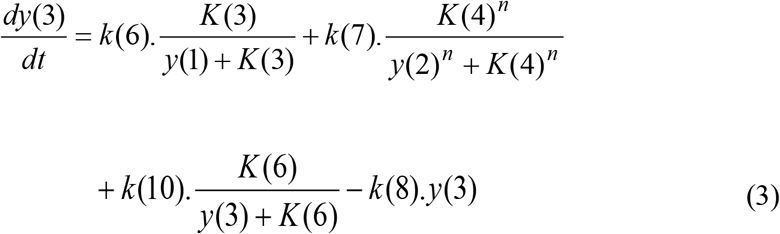

All parameters are described in Table 1.

### Systems modeling and determination of successful memory

The time evolution of the gene expression levels of *y*(1), *y*(2) and *y*(3) were simulated both deterministically and stochastically. For a deterministic systems description, the ordinary differential equations (Eqs. 1-3) were solved in MATLAB (Mathworks) with standard solvers. Stochastic trajectories were simulated with the Gillespie algorithm [63].

In all model simulations, the deterministic and stochastic time courses of *y*(1), *y*(2) and *y*(3) were simulated for 1,000 simulation time steps. The input signal *S* was applied from simulation step 250 to 500. Deterministic memory was defined as sustained protein levels in the numerical integration of the rate equations during the subsequent period from simulation steps 500 to 1,000 after stop of the signal *S*. Similarly, stochastic memory was assessed as sustained protein levels in the stochastic simulations in the period from simulation steps 500 to 1000 after stop of the signal *S*. While arbitrary, a threshold of 1000 simulation time steps yielded a robust and readily accessible criterium to assess memory. Due to the probabilistic nature of the stochastic simulations, robust stochastic memory for a given set of parameters was defined as successful memory if 18 out of 20 stochastic simulations, i.e. 90%, yielded persistent memory [12].

### Theoretical model comparisons

All parameters were set as to render the MRN and MRN-NA models as comparable as possible (Tables 1, 2) [12, 64]. With the exception of the additional negative autoregulation loops, this meant using the same parameters throughout. The parameters of the negative autoregulation were tuned to conserve the steady state levels of *y*(2) and *y*(3) between the MRN and MRN-NA models (Tables 1, 2). Indeed, the high steady-state levels of both the models were set to the same, while the low steady-state levels of them could be as similar as possible. Note that it is impossible to set the low steady-state levels to the same. Given the asymmetry of the models (activation of *y*(2) and suppression of *y*(3) by *S*), the deterministic steady-state levels of *y*(2) and *y*(3) always show opposing behavior: when the steady-state level of *y*(2) is high, that of *y*(3) is low and vice versa. Our parameters choices conserved both the low and high steady states of *y*(2) and *y*(3) between the MRN and MRN-NA models (Texts S1, S2, [12])

To systematically identify parameter choices that can give rise to successful memory, phase diagrams of the memory region as a function of the two most critical parameters, the Hill coefficient and dissociation constants in the mutual repression cycle, were computed by successive simulation at different parameter combinations.

Intrinsic noise in gene expression was quantified by computing the coefficient of variation (CV) of the levels of *y*(2) and *y*(3) during the input signal *S* from the stochastic simulations. To eliminate transition effects, we considered the period from simulation step 270 to 500, i.e. omitting the first 20 simulation steps of the signal period (Fig. 2F). In the same manner, we estimated the CVs of the levels of *y*(2) and *y*(3) after stop of the signal, for the period from simulation step 500 to 750 (Fig. 2G).

### Stochastic potential and probability density function

The reaction rate equations (Eqs. (1–3)) were converted into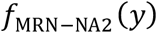 and 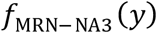(Text S2). Using the quasi-steady-state assumption, we solved the probability density at the steady state (at *t* → ∞). At *t* → ∞ *y*(2) and *y*(3) definitely approach to the steady state, i.e., 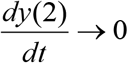 and 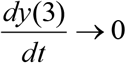. Therefore, we assumed 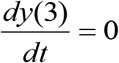 to solve the ODE of 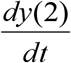 at *t* → ∞. In a similar manner, we assumed 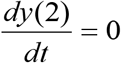 to solve the ODE of 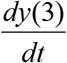 at *t* → ∞.

Here, we illustrated how a one-variable equation 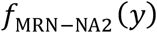 in a stochastic environment is given by:

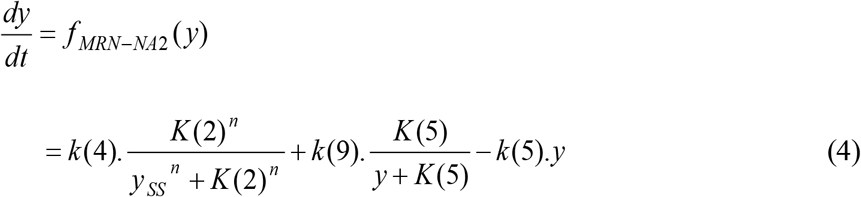

where

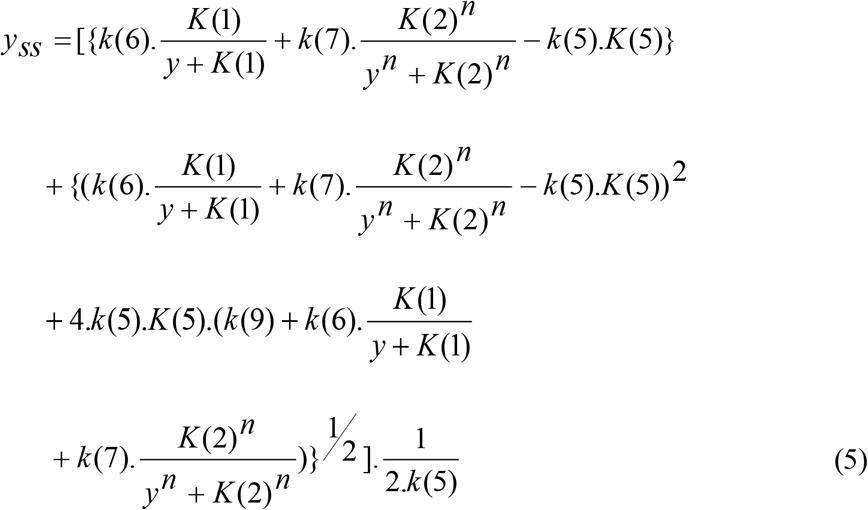

This equation can be described by the birth-and-death stochastic processes [12, 21, 48, 65]:

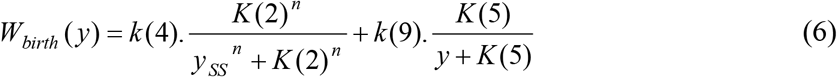

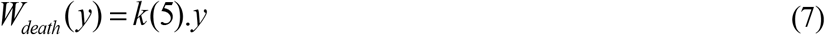

where *y*_*ss*_ is given by Eq. (5)

The corresponding chemical master equation was given by:

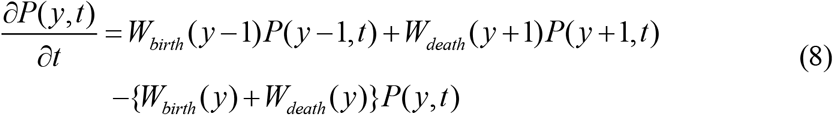

where *P*(*y, t*) was the probability density of protein concentration *y*. Next, the chemical master equation was transformed into the Fokker-Planck equation [12, 21, 49, 65-67]:

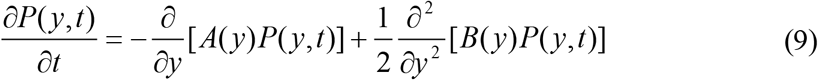

where

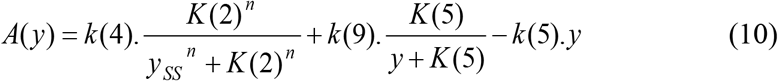

and *y*_*ss*_ is given by Eq. (5). The noise function is given by:

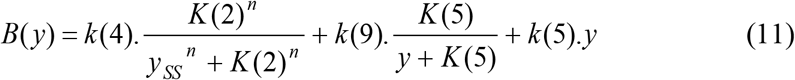

where *y*_*ss*_ is given by Eq. (5). In the same manner, the Fokker-Planck equations of the four one-variable equations including 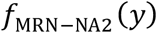 were solved under the following conditions (Text S2):

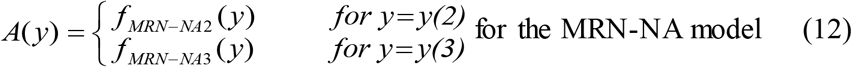

and the noise functions were given by:

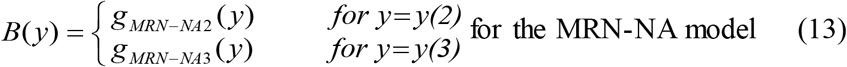

Finally, we consider the stochastic potential analysis. The limit of *P*(*y, t*) at *t* → ∞ yields *P*_st_(*y*), the stationary probability density function of *y* [12, 21, 58, 65], which is given by:

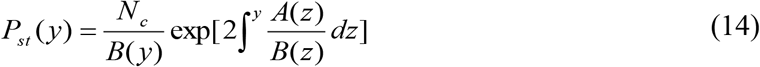

where *N*_*c*_ is the normalization constant. Eq. (14) can be recast in the form:

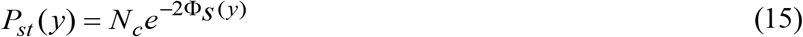

where

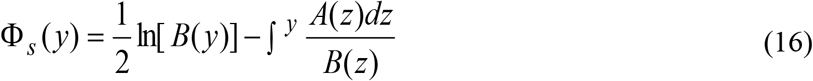

is called the stochastic potential of *f*(*y*) [56, 57, 65].

### Mean first-passage time analysis

In gene expression, the stability of a steady state has to be estimated in the presence of noise. The stability of a steady state of a stochastic system can be estimated by the mean first-passage time (MFPT), which describes the expected time within which the system leaves a stable steady state due to random fluctuations. An equilibrium point can exit from its minimum potential due to the effect of noise. The exit time depends on the specific realization of the random process; this is known as the first passage time. The MFPT is the average of the first passage times over many realizations. In the context of anticipating phase shifts, the MFPT provides a useful tool to characterize the time - scale on which a phase transition is likely to occur.

Let us consider 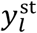 and 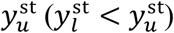 as two steady states corresponding to a low and a high protein concentration, respectively, separated by the unstable steady state defining the potential barrier 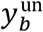 (i.e., the unstable equilibrium point). The basin of attraction of the state 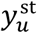 extends from 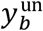 to +∞, as it is to the right of 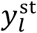. Let *T*(*y*) be the MFPT to state 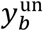 starting at 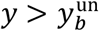. *T*(*y*) satisfies the following ordinary differential equation [21, 58, 66, 67]:

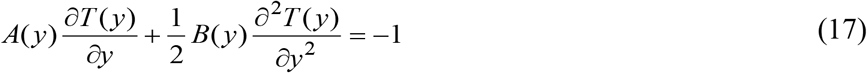

with boundary conditions:

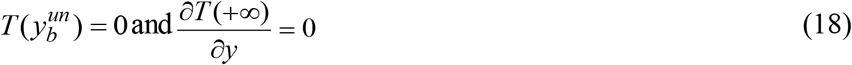

By solving Eqs. (17–18), the MFPTs of 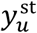 and 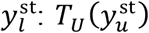 and 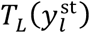 are calculated to state 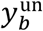 for the basin of attraction of the state 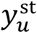 extending from 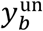 to +∞ and for the basin of attraction of the state 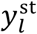 which extends from 0 to 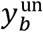, respectively, as follows:

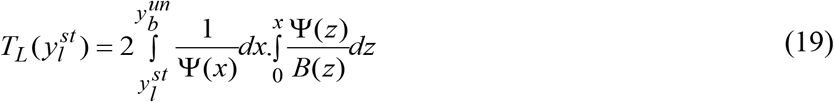

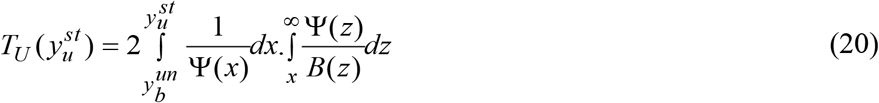

where

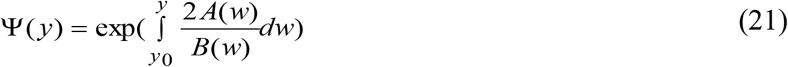

with *y*_0_ = 0 for the 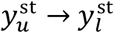 transition and 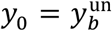 for the 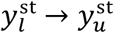 transition. A high value of the MFPT of a steady state protein level indicates that the level is sustained for a longer time, whereas a low value indicates that the protein can readily leave the steady state and quickly transition to another level.

## Acknowledgments

We are grateful to Dr. Kazuhiro Maeda for helpful discussions. SP holds the Canada Research Chair in Computational Systems Biology and acknowledges funding from the Merck Sharp Dohme program of the Faculty of Medicine of the Université de Montréal.

## Code availability

The computer code used to perform all analyses is available at https://github.com/ABMSUH/MRN-and-MRN-NA-models

## Author Contributions

Conceptualized the research: ABMSUH, HK; Developed all methodology and performed all analysis: ABMSUH; Interpreted the results: ABMSUH, HK, SP; Wrote the original draft: ABMSUH; Reviewed and edited the final manuscript: ABMSUH, HK, SP.

## Competing Interests

The authors declare that they have no competing interests.

## Notes

#### Summary of Updates

Revised introduction; reduction and consolidation of figures and results sections to better differentiate what is new and what was recalculated for comparison to a previously published paper.

